# *ξ-π*: a nonparametric model for neural power spectra decomposition

**DOI:** 10.1101/2023.12.03.569765

**Authors:** Shiang Hu, Zhihao Zhang, Xiaochu Zhang, Xiaopei Wu, Pedro A. Valdes-Sosa

## Abstract

The power spectra estimated from the brain recordings are the mixed representation of aperiodic transient activity and periodic oscillations, i.e., aperiodic component (AC) and periodic component (PC). Quantitative neurophysiology requires precise decomposition preceding parameterizing each component. However, the shape, statistical distribution, scale, and mixing mechanism of AC and PCs are unclear, challenging the effectiveness of current popular parametric models such as FOOOF, IRASA, BOSC, etc. Here, *ξ*-*π* was proposed to decompose the neural spectra by embedding the nonparametric spectra estimation with penalized Whittle likelihood and the shape language modeling into the expectation maximization framework. *ξ*-*π* was validated on the synthesized spectra with loss statistics and on the sleep EEG and the large sample iEEG with evaluation metrics and neurophysiological evidence. Compared to FOOOF, both the simulation presenting shape irregularities and the batch simulation with multiple isolated peaks indicated that *ξ*-*π* improved the fit of AC and PCs with less loss and higher F1-score in recognizing the centering frequencies and the number of peaks; the sleep EEG revealed that *ξ*-*π* produced more distinguishable AC exponents and improved the sleep state classification accuracy; the iEEG showed that *ξ*-*π* approached the clinical findings in peak discovery. Overall, *ξ*-*π* offered good performance in the spectra decomposition, which allows flexible parameterization using descriptive statistics or kernel functions. *ξ*-*π* may be a promising tool for brain signal decoding in fields such as cognitive neuroscience, brain-computer interface, neurofeedback, and brain diseases.

## I. Introduction

**N**EURAL oscillations are the repetitive or periodic activities centering frequencies, shaping the neuron extracellular field potentials and transmitting activities between neural masses. Aperiodic and transient neural activities are ubiquitously produced by the neural assemblies. The synchronous activities by millions of neurons are macroscopically detectable by intracranial Electroencephalography (iEEG), EEG, and other brain recordings [1]. Although the neural underpinnings and interpretation of periodic and aperiodic activity are significantly different, the two activities constantly mix in brain recording producing two typical shapes in the neural power spectra. The 1/f-like background spectra are the aperiodic component (AC) attributed to the aperiodic activity, while the spectral peaks are the periodic component (PC) reflecting the periodic activities. Studying neural activities has to quantitatively characterize the AC and PCs, such as the AC offset and slope and PC center frequency (CF), bandwidth, and amplitude [2]. AC has potential applications in studying brain development [3]–[5], cognitive performance [6], and psychiatric disorders, such as attention deficit hyperactivity disorder [7] and schizophrenia [8]. The studies on PCs can help reveal the nature of the cognitive process and the origin of brain dysfunction [9]–[12]. The PC parameters are closely related to brain functions, such as the vision using the alpha rhythms [13], the motion with mu rhythms [14], the attention using the gamma rhythms, the cognitive processes [1] and the pathological state [15], and brain aging [5], [16]. Nevertheless, the mixing of AC and PCs hinders the study of brain functions correctly, requiring a precise and robust spectral decomposition model.

Several models have been proposed for EEG/MEG spectral decomposition, as shown in Table I. They can be distinguished by the scale in which the spectra are fit: natural or logarithmic. *ξ*-*α* model adopts the Student’s *t* curve function to fit AC and PCs in the natural scale, centering attention on a sole alpha peak [17]. BOSC (Better OSCillation detection) detects the PCs based on a chi-squared distribution and linearly fits the AC in the log-log space assuming a 1/f form [18]. IRASA (Irregular Resampling Auto Spectral Analysis) can estimate AC according to the different robustness of fractal and periodic activities to resampling but cannot isolate the peaks [19]. As a recent and popular model, FOOOF (Fitting Oscillations & One Over f) employs the Gaussian and the power law functions to fit the PCs and AC in the log scale, respectively [20]. Further, adapted from FOOOF, the SPRiNT (Spectral Parameterization Resolved in Time) allows for the time-resolved spectral decomposition [21], and the PAPTO (Periodic*/*Aperiodic Parametrization of Transient Oscillations) considers the spectral peak as arising from transient events [22]. These models may be limited to incomplete decomposition, parametric fitting with hard kernels, or log scale transformation.

**TABLE I.** Summary of spectra decomposition models

| Models | Spectra decomposition |  | Statistical distribution |  | Fitting scale | Mixing mechanism |
| --- | --- | --- | --- | --- | --- | --- |
|  | AC | PC | AC | PC |  |  |
| ξ-α | ✓ | × | T curve | T curve | Natural | Additive |
| BOSC | ✓ | × | Power law | Chi-square | Log-log | Additive |
| IRASA | ✓ | × | Power law | - | Log-log | Additive |
| FOOOF, PAPTO, SPRiNT | ✓ | ✓ | Power law | Gaussian | Log power | Multiplicative |
| ξ-π | ✓ | ✓ | **Monotonicity** | **Unimodality** | **Natural** | **Additive** |

Although the shape, statistical distribution, scale, and mixing mechanism of background and periodic spectral components are not yet clearly understood, opinions can be drawn from practices. Firstly, as shown in Fig. 1, the iEEG spectral shape may be of greater diversity and present more peaks than the scalp EEG/MEG spectra. The shapes of AC and PC may not strictly follow the power law function and symmetry bell shape, respectively. The left and right petals of a spectral peak may be convex to CF. Two neighboring peaks easily overlap if their CF are close, where the parametric models may mistakenly fit the two as one peak. Prior studies found that the AC may be either scale-free activity or pink noise, complying with the power law distribution [23]. Whereas the physical nature of 1/f like AC is not yet determined, which may be modeled from the 1-order autoregressive process, the point process, the sum of Lorentzians, or the avalanches in self-organized criticality [24]–[26]. Thus, the shape of AC and PC may be irregular, with the exact statistical distribution remaining unknown. Secondly, the existing models are inconsistent in scale transformation. *ξ*-*α* parametrizes the spectra in the linear power vs. frequency space, BOSC detects the AC in the log-power vs. log-frequency space, IRASA estimates the fractal component of power spectra in the linear power vs. frequency space but fits the power law in the log-power vs. log-frequency space, while FOOOF, SPRiNT, and PAPTO decompose the spectra in the log-power vs. frequency space. Inconsistencies of scale transformation imply two fundamentally different mixing mechanisms of AC and PC: additive and multiplicative modulations. The additive model is the most parsimonious viable for decoding the resting brain activity. The level of 1/f activity and alpha power are not positively correlated in line with the additive but not the multiplicative model [27]. One should not apply log or other nonlinear transformations to the power and frequency before decomposition, avoiding under- or over-estimating the parameters [28]. Overall, any bias to one component introduced in parametric decomposition and nonlinear transformation will contaminate the fit of other components. Misfit spectral components will derive biased oscillatory parameters. This is critical to practical inferences and theoretical interpretations in quantitative neurophysiology.

**Fig. 1.**
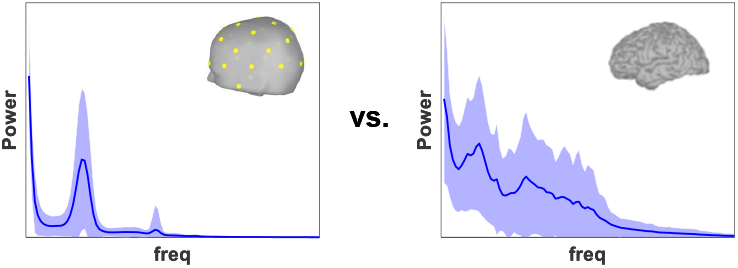
Comparison between 130 channels of scalp EEG spectra (left) and 123 channels of iEEG spectra (right).

To address the known issues, we designed a nonparametric decomposition model following the natural scale and additive mechanism, allowing for parameterization as a subsequent step. Since nonparametric models do not need to predetermine a function but learn from data, they help fit the natural shape characteristics, i.e. monotonicity for AC and unimodality for PC. Several models may suit this problem, such as the shape regression, the generalized additive models for location, scale, and shape, the unimodal smoothing, and the shape-constrained additive models [29]–[32]. Besides, the unmixing step isolates unknown components from power spectra. From the view of Bayesian statistics, this step estimates the latent population variances from observed sample variances, which may be properly resolved in the expectation-maximization framework.

## II. Methods

### A. Mathematical basis of ξ-π

#### 1) Decompositon

The power spectra estimate commonly takes the smoothed periodogram or multitapers method by segmenting the recordings into quasi-stationary epochs that may be segments, windows, or tapers elsewhere. Assuming the FFT size equals *N*, the sampling rate is *fs*, the frequency resolution is Δ*f* = *fs/N*. Note that the complex normal distribution of Fourier coefficients is asymptotically independent across frequencies due to the stochastic properties of finite Fourier transforms [33], [34]. Thus for the segment *s* = 1, …, *n*_*s*_ and the frequency *f* = 1,…, *n*_*f*_, the Fourier coefficient *y*_*f,s*_ ∈ ℂ^1^ is additive over multiple processes/components *k* = 1,…, *n*_*k*_ of brain activities, which is expressed as

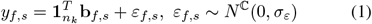

where **1**_*n*_ ∈ ℝ^*n×*1^ is a vector of ones, the Fourier coefficients corresponding to multiple brain processes follow a multivariate complex normal distribution 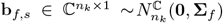, the model error is assumed with the constant variance across frequencies, the diagonals of matrix **Σ**_*f*_ are 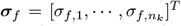, and the off-diagonal entries represent the cross spectra between two different brain processes.

Estimating the population variance of Eq. 1 derives the mix of variance components, i.e. the mixed power spectrum

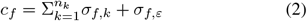

where *σ*_*f,k*_ ∈ ℝ^1^ is the power spectrum of the component *k* at the frequency *f*.

The observed power spectrum *p*_*f*_ is the sample variance over finite segments, which approximates to *c*_*f*_

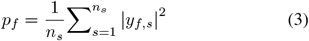

The spectral decomposition is to estimate the variance *σ*_*f,k*_ of latent variables **b**_*f,s*_ from *p*_*f*_. This is solved by the expectation-maximization (EM) algorithm. The E and M steps take the effect of ‘re-decompose’ and ‘individual fit’ accordingly as shown in Fig. 2. The initial condition is defined as 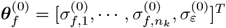 where the 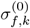 is generated by the Student’s *t* function [17] as shown in Fig. 2(c), and the 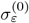 was set to 0.01 for all frequencies.

**Fig. 2.**
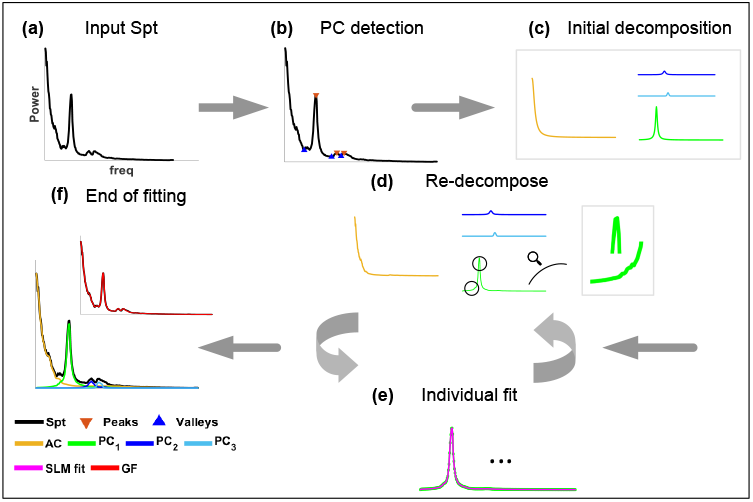
Schematic flowchart of *ξ*-*π*. **(a)**. input spectra; **(b)**. peak detection; **(c)**. initial decomposition using the Student’s ***t*** function; **(d)**. re-decompose spectra through E step; **(e)**. fit individual components using SLM through M step; **(f)**. output the AC and PCs at the end of fitting. **Spt**: Spectra, **GF**: global fit.

In the E step, given *i* the number of iterations, the parameter 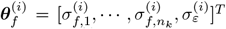 and the pseudo mixed power spectrum 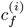 produced in the M step, taking minimum norm least square solution to estimate latent variables **b**_*f,s*_

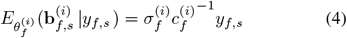

which shows that the Fourier coefficient decomposition behaves as Wiener filtering.

In addition, the pseudo power spectra of each component and model error are

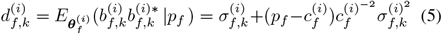

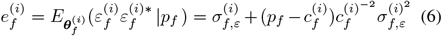

And utilizing the variance components model, the complete negative log-likelihood is

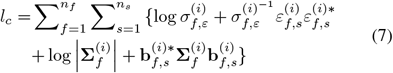

In the M step, given the pseudo power spectra 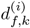 and 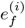, the *Q* function is

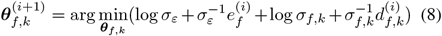

The solution to this convex problem is the estimation of spectra component 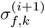 and decomposition error 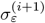 Note that the two parts in this *Q* function follow an identical structure known as the Whittle likelihood [33]–[35]. Theoretically, the Whittle likelihood holds statistical consistency to the spectra density estimation [36], [37]. The Whittle likelihood [34] of the spectral component is

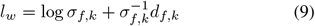

This allows estimating multiple spectral components in a non-parametric way by incorporating the shape constraint detailed in the section II-A.2. The spectra decomposition error for all frequencies is estimated by taking average reference [38].

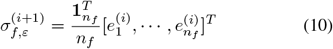

#### 2) Nonparametric fitting

The estimation of 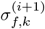 in the M step is solved by the shape language modeling (SLM) based on the cubic spline and shape priors [39], [40]. The priors from the natural shape characteristics are monotonically decreasing for AC and rising first to CF then falling for PC. The SLM lowers uncontrollable flexibility of spline fitting by prescribing shape priors into a set of shape primitives, performing better fit than parametric models and nonlinear regression in a shorter time. Here, the strategy behind SLM is in line with the penalized Whittle likelihood because both are Bayesian approaches. In the M step, the prescription is piecewise polynomial Hermite interpolation jointing with different knots. Specifically, for AC fit, the number of knots is set as *n*_*f*_ */*4, and the shape is set as monotonous decreasing; for PC fit, the number of knots is set as *n*_*f*_ */*3, and the shape is set as a simple peak.

### B. Algorithm implementation

As shown in Fig. 2, the implementation of *ξ*-*π* has the following steps:

a. **Input spectra (Spt)**: Dividing the spectral values by their maximum to benefit the generalizability of parameters set for peak detection. The normalization removes the effect of absolute magnitude and judges the peaks by the scale-free shape properties.
b. **PC detection**: The PC detection is to isolate the prominent peaks forming the concave shapes, which needs to localize the spectral peaks and valleys. The MATLAB ‘findpeaks’ function is efficient in detecting peaks. The valleys are detected from the up-down inverted spectra, i.e., the negative spectra. The parameters were set to:’min peakwidth, 0.9’, ‘min peakheight, 0.05’, and’ min peakprominence, 0.025’.
c. **Initial decomposition**: Based on the leftmost maximum and the first valley right near the maximum, the Student’s *t* function was used to initially fit AC. Then it sequentially took the next peak-valley pair to initially estimate PCs.
d. **Re-decompose**: All initialized components were submitted to the EM algorithm, where the E step regenerates pseudo power spectra of individual components (Eq. 4 5 6).
e. **Individual fit**: The M step fits the individual component using SLM. The (d-e) loop will be executed iteratively to minimize the incomplete negative log-likelihood.
f. **End of fitting**: The iteration ends when the EM algorithm iterates to a preset number of times or the incomplete negative log-likelihood is lower than the threshold.

### C. Parameterization

It is necessary to parameterize the AC/PC to quantify brain rhythms and represent neural dynamics. *ξ*-*π*, as a non-parametric model, allowing flexible parameterization using user-preferred models on actual needs. For the later model comparison and evaluation, the same function as FOOOF was adopted to quantify decomposed AC/PCs by *ξ*-*π*, that is

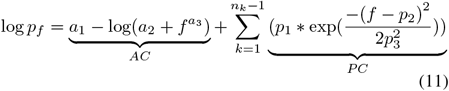

where *a*_1_,*a*_2_, and *a*_3_ are the AC offset, AC knee, and AC exponent, respectively. *a*_2_ = 0 if the AC knee is not accounted for. In addition, PC is modeled by the Gaussian function in the Eq. 11 where *p*_1_,*p*_2_ and *p*_3_ stand for the maximum power, CF, and half of the bandwidth in the log scale. Note that the scale transformation is made for comparison.

### D. Simulation

Prior findings show that AC commonly exists over the global brain but PC may merely exist over local brain regions [5], [41], [42]. The time series and its spectra of the aperiodic activity or the periodic oscillation with sole spectra peak can be simulated, providing an objective standard to assess how *ξ*-*π* behaves. The general scheme is synthesizing the AC and PCs separately and then summing them as mixed spectra. The PC simulation follows two different ways: (1)”PC-resampling”: pruning band-limited periodic spectra with shape irregularities after the AC was estimated by IRASA and removed from real spectra; (2)”PC-sine waves”: estimating the spectra from time series composed by a few sine waves that differ in CFs for batch simulating spectra with multiple isolated peaks.

#### 1) AC

Either parametric or nonparametric models subjectively used to simulate AC will later show favor to the corresponding decomposition model in the validation process. A theoretically valid approach is learning the AC prototypes from real spectra by unsupervised learning. Thus, we empirically and randomly selected a channel from the no-peak set as the ground truth AC as shown in Fig. 3B. The no-peak set consisting of 297 channels was clustered from the iEEG database detailed in the section II-E.2 [43].

**Fig. 3.**
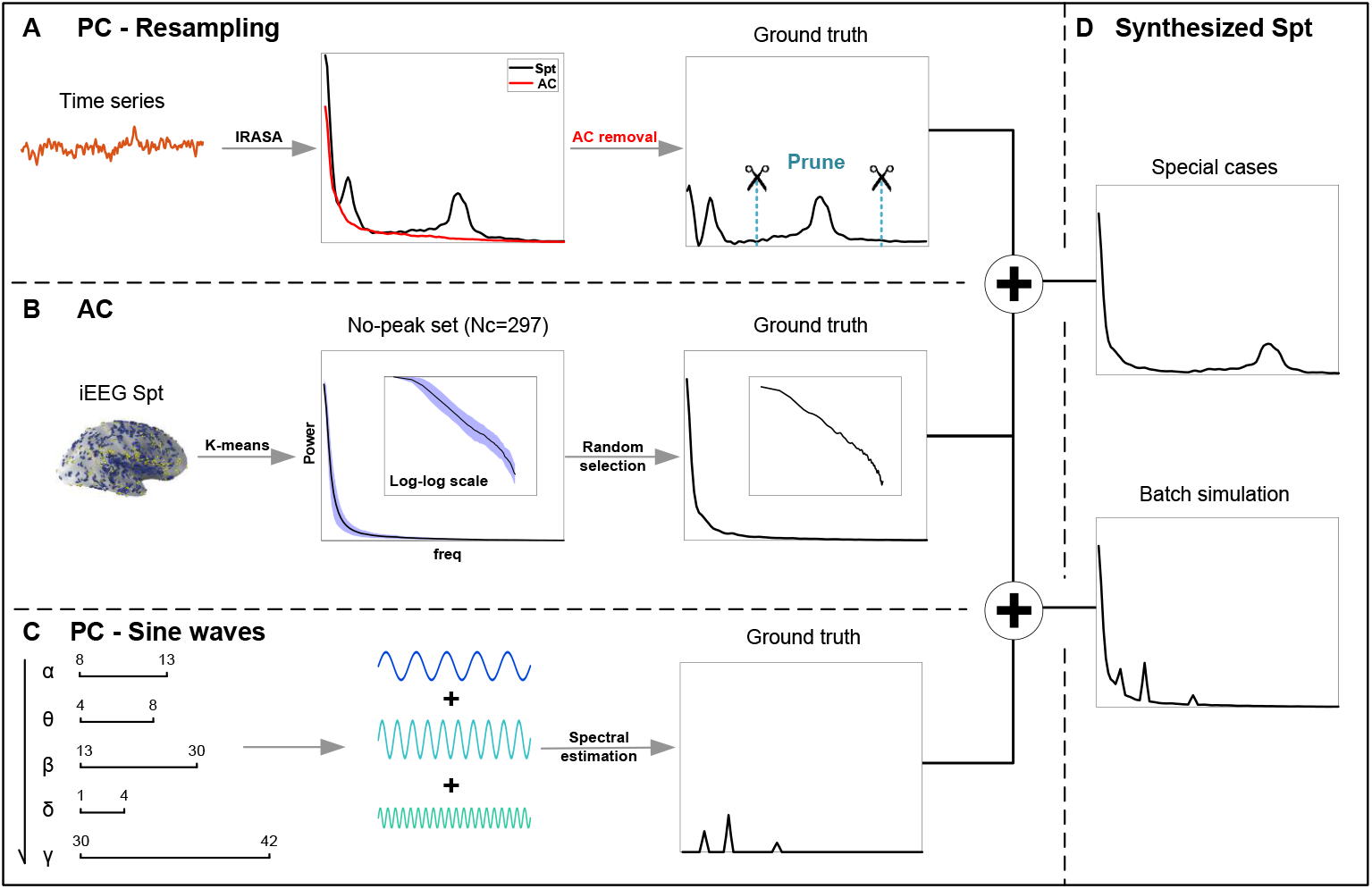
The spectra simulation pipeline. **(A)**. PC-Resampling: IRASA was applied to remove AC; then the skewed, overlapping, or concave upward peak was pruned from mixed PC; **(B)**. AC was randomly selected from the no-peak set (297 channels); **(C)**. PC-Sine waves: mixing no more than 5 types of sine waves with CFs in different narrow bands; the priority of sine waves follows the order [*α, θ, β, δ, γ*]; the mixed PC was obtained after estimating the spectra of mixed time series; **(D)**. Synthesized Spt: Special cases – adding the PC from (A) to the AC; Batch simulation: adding the PC from (C) to the AC.

#### 2) PC-Resampling

As shown in Fig. 3D, the simulation of special cases aims to validate the decomposition effects on the spectra that present a peculiar peak. Although, commonly, the spectra peak does not strictly follow the symmetric bell shape, there is no straightforward way to simulate the peculiar peak by generating the time series first. Alternatively, the simulation was done by learning from real iEEG activity, as shown in Fig. 3A. Firstly, IRASA was applied on the iEEG channels in the precentral gyrus to estimate AC; secondly, the mixed PC was isolated by subtracting AC from the original spectra; thirdly, the PC with shape irregularity was obtained by pruning the lower and higher frequencies. Finally, the three special cases of PCs were simulated: (a) **overlapping peak**: the CFs of two peaks were close so that the left slope of the peak with higher CF rode on the right slope of the peak with lower CF, forming a high amplitude valley between the two peaks. (b) **skewed peak**: the left and right slopes of a peak were asymmetric around its CF. The long tail on one side of the peak slopes made the peak skew toward the other side. (c) **concave upward peak**: the left and right slopes of a peak were concave upward, forming a sharp hat on the top.

#### 3) PC-Sine waves

Large samples of spectra were generated through batch simulation for loss statistics and quantitative comparison, as shown in Fig. 3C. The number of peaks takes the range of 0-5. The CFs are required to distribute in different narrow bands and follow the priority of *α*(8-13Hz), *θ*(4-8Hz), *β*(13-30Hz), *δ*(1-4Hz), *γ*(30-42Hz). The priority of narrowband peak comes from the empirical probability of the peak occurrence in a specific band. In addition, two peaks are avoided from overlapping by setting two CFs distant at least 3Hz. For instance, when simulating a PC with 2 peaks, we randomly set the CFs, one in the *α* band and the other one in the *θ* band. Besides, the amplitudes of sine waves increase with the CFs shifting from *δ, θ* to *α* and decrease with the CFs shifting from *α, β* to *γ*. Finally, the 100 samples by 60 seconds of time series were generated by the repetitive simulation.

#### 4) Loss Statistics

##### Mean squared error (MSE)

The MSE of decomposed AC is expressed as:

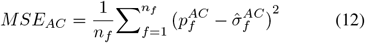

where 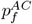 is the simulated ground truth AC and 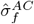 is the decomposed/fitted AC. The MSE of decomposed PC takes the form:

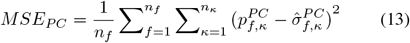

where the 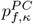 is the simulated ground truth of PC, 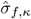 is the decomposed/fitted PC, and *κ* is indexing the peaks.

##### Number of peaks

The set number of peaks in batch simulation is taken as the ground truth to check the decomposition effects in PC detection. The number of decomposed PCs was counted and compared with the simulated truth. The overestimation ratio (OER) and the underestimation ratio (UER) are defined as

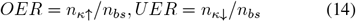

where *n*_*bs*_ denotes the total number of batch simulated samples, and *n*_*κ↑*_(*n*_*κ↓*_) indicates the number of samples that detected more (less) peaks than the set.

##### Identification of CFs

To quantify the PC detection, the frequency bins of each simulated sample are labeled as ‘CF’ or ‘no CF’ based on whether the frequency was set as a CF or not. Thus, being CFs or not across the frequencies and the repeated samples form the binary discrimination problem, allowing for the use of model evaluation metrics such as accuracy, precision, recall, and the F1-score.

### E. Real data and analysis

#### 1) Sleep state classification using CCSHS EEG

The open Cleveland Children’s Sleep and Health Study (CCSHS) [44], [45] database contains the polysomnography recordings from 515 teenagers while the subject was sleeping. Each subject underwent the recording of two electrodes, C3 and C4, lasting for 41220s. The data were segmented into epochs with 30s duration, inspected to manually remove artifacts, and annotated as Wake, NREM-1(N1), NREM-2(N2), NREM-3(N3), and REM states by trained technicians. Here, the N2 was used to represent the NREM state because of its overwhelming dominance compared with the N1 and N3, both of which have short coverage and low occurrence [46]. The CCSHS EEG was 0.5-50Hz bandpass filtered.

For each sleep state, the C3 and C4 epochs from randomly picked two subjects were concatenated into 10 mins recordings by ignoring the differences across electrodes and subjects. Note that the data concatenation aims to later perform both time-varying spectral decomposition and cross-subject/electrode sleep state prediction. Since the whole sleep recording is long enough, we repeated this procedure sequentially in the time order without using the same epoch twice to create multiple data collections. The EEG recordings of 3 states by 10 mins by 19 collections were prepared for sleep state classification. Hence, we finally conducted 19 times of independent sleep state classification experiments.

For the recording per state per collection, the time-varying spectra were estimated by applying a 30s sliding window with 50% overlapping, which yielded 39 windows. The time-varying AC parameters, exponent and offset, were obtained after parameterization. Therefore, we obtained 39 pairs of time-varying AC parameters per state per experiment. Later, the linear discriminant analysis (LDA) was performed to predict sleep state based on the AC exponents [47]. For cross-validation, we used the leave-one-out strategy which partitioned the 39 windows as a training set of 38 windows and a test set of 1 window and repeated 39 folds. Three LDA predictors were constructed for differentiating the states of Wake-N2, Wake-REM, and N2-REM, respectively. The distinguishability of AC exponents across states was further evaluated through prediction accuracies.

#### 2) Peak discovery using MNI iEEG

The open MNI iEEG dataset was collected from 106 epilepsy patients (33.1±10.8 yrs.) over multi-sites and curated by the Montreal Neurological Institute and Hospital (MNI) [43]. In addition to epileptic lesion regions, the healthy brain regions of individual patients underwent the iEEG recordings as well for brain state monitoring and clinical evaluation. Ethical approval was granted at the MNI (REB vote: MUHC-15-950). Although the number of iEEG channels planted into the healthy brain regions was different for each patient, the 1772 channels accumulated from 106 patients broadly covered the whole brain in the MNI space, providing a spectral atlas of the human brain. The sampling rate was downsampled from the raw 1000Hz to 200Hz. A 64s segment was selected from each channel, which contains no epileptic activity and other artifacts. With the 1772 channels of iEEG, the decomposition models were validated from the following metrics:

##### WLS

The weighted least square (WLS) measures the residuals between the real spectra and the reconstructed global fit (GF) summing over the decomposed AC and PCs. The smaller the WLS, the better the model. The WLS is expressed as

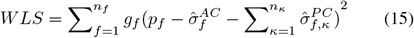

where *p*_*f*_ is the real spectra, 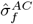 and 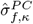 are the decomposed AC and PCs, and *g*_*f*_ is a normalized weight vector with relatively large values in the *α, β* and *θ* bands. The spectral magnitudes are considerably different between low and high frequencies, resulting from the typical characteristic of exponentially decaying in magnitude across broadband. The *g*_*f*_ was designed as an elastic ruler to improve the reliability of the residual calculation.

***R***^**2**^. The *R*^2^ is calculated by taking the square of the correlation coefficient between GF and Spt. It indicates the similarity between the GF and the original spectra.

##### Whittle likelihood

The Whittle likelihood as Eq. 9 takes the form of negative log-likelihood which is a statistical measure to indicate how the model is close to the data. Thus, the smaller the Whittle Likelihood, the better the model.

##### Peak discovery proportion

The spectra decomposition discovered the peaks and allowed counting the proportion of the number of channels presenting a peak within a narrowband to the total channel number in a brain region. The study [43] reported that four brain regions presented the specific peak discovery proportions after inspection. That is, 72% of 36 channels in the hippocampus have a delta peak at ∼1Hz; 68% of 19 channels in the cuneus have an alpha peak at ∼8Hz; 72% of 39 channels in the opercular part of the inferior frontal gyrus (OFG) have a beta peak at 20-24Hz; 64% of 123 channels in the precentral gyrus has a beta peak at 20-24Hz. These findings are taken as the empirical standard to assess the decomposition effects on peak detection. After spectra decomposition, the peak discovery proportion was counted for these regions and visually validated for each channel.

## III. Results

### A. Simulation Analysis

#### 1) Special Cases with Shape Irregularity

Fig. 4 is the demonstration of how *ξ*-*π* performed on three typical spectra with peculiar shapes. **As to “overlapping”** in Fig. 4A, though both models found 1 AC and 2 PCs, *ξ*-*π* enabled fitting the AC and 2 PCs with good coincidence and alignment to the original Spt and the CFs, while FOOOF underfitted the AC and the overlapping peak with a major peak and a minor peak which is just the low amplitude tail in the original Spt. The AC and PC fit show clearly that *ξ*-*π* can fit the original Spt with high coincidence, but FOOOF fitted the AC much lower than the spectrum at a very low frequency and failed to fit the overlapping peak by mixing two Gaussians.

**Fig. 4.**
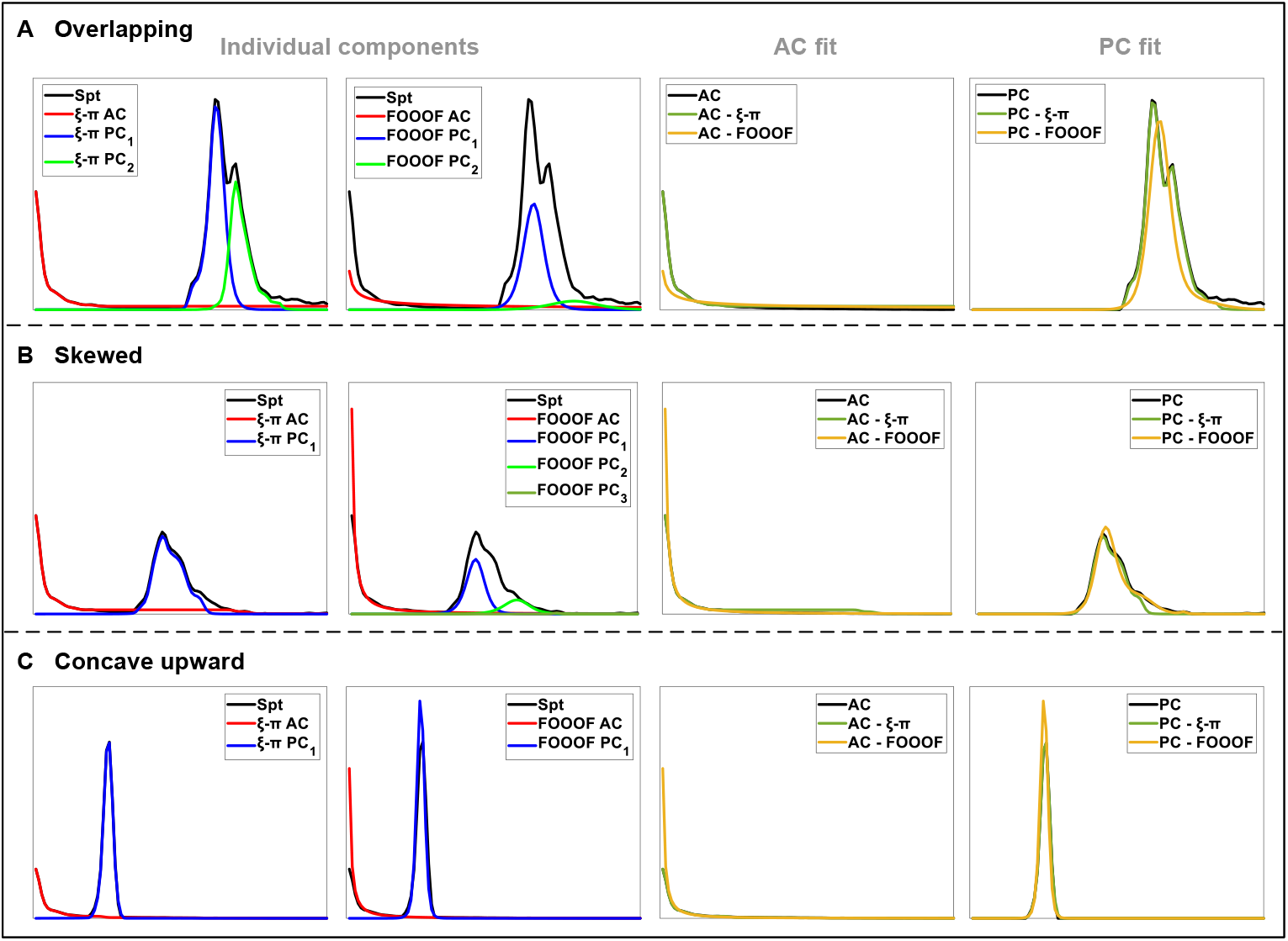
Special cases with shape irregularity. **(A)**. overlapping peak; **(B)**. skewed peak; **(C)**. concave upward peak. The 1-2 columns display the decomposed individual components. The 3-4 columns are the AC fit to the ground truth AC and the composite PC fit to the ground truth PC by summing over the individual PCs. Spt: simulated spectra; All the plots are the power against the frequency(Hz) in the natural scale.

**In terms of “skewed”** in the Fig. 4B, *ξ*-*π* decomposed Spt into 1 AC with good coincidence and 1 PC with the low amplitude long tail being underfitted, but FOOOF failed to fit the AC at very low frequency and fitted the sole skewed peak with two peaks. The AC fit shows that the AC decomposed by *ξ*-*π* may not exponentially decay to zeros at high frequencies, and FOOOF fitted much higher than the spectrum at very low frequencies. The PC fit displays that *ξ*-*π* generally coincided with the ground truth PC except for the long tail with low amplitude and high frequencies, but FOOOF fitted higher amplitude than the ground truth peak, shifted the CFs toward higher frequency, and did not fit the right slope of the PC.

**Regarding “concave upward”** in Fig. 4C, *ξ*-*π* enabled to fit the original shape of ground truth AC and PC, but FOOOF overfitted the AC at very low frequencies and decayed fast to zeros, although the ground truth AC are non-zeros, and FOOOF failed to fit the ground truth PC with high peak amplitude and shifted CF toward lower frequency.

Thus, *ξ*-*π* was good at keeping the natural shape characteristics, although the shape may be extraordinary, whereas FOOOF may bring about extremely high amplitude in very low frequency, decay exponentially to zeros in AC fit, and fail to fit the peak shape even shifted the CFs in the PC fit.

#### 2) Batch Simulation and Loss Statistics

Fig. 5A displays the distribution of set CFs in batch simulation. For each row, the grids in color are the set CFs in a simulated sample, and the number of grids is the same as the number of simulated peaks. The number of peaks and the distribution of the CFs were different for each simulated sample. Fig. 5B compares the peak number between the simulation and the fit. As to columns from left to right, the preset number of peaks is [0, 1, 2, 3, 4, 5]. The number of simulated samples with a preset peak number was [19, 17, 17, 11, 18, 18] correspondingly, which equaled the sum of bubble sizes in the same color within a column. The bubbles along, above, and below the diagonals indicate the proper estimation, the overestimation, and the underestimation of the peak number, respectively. The OER (UER) of *ξ*-*π* and FOOOF were 0% (7%) and 47% (30%). This means that *ξ*-*π* is reliable and not easy to overestimate and underestimate the peak number; FOOOF may not only easily detect more peaks but also neglect more peaks than *ξ*-*π*.

**Fig. 5.**
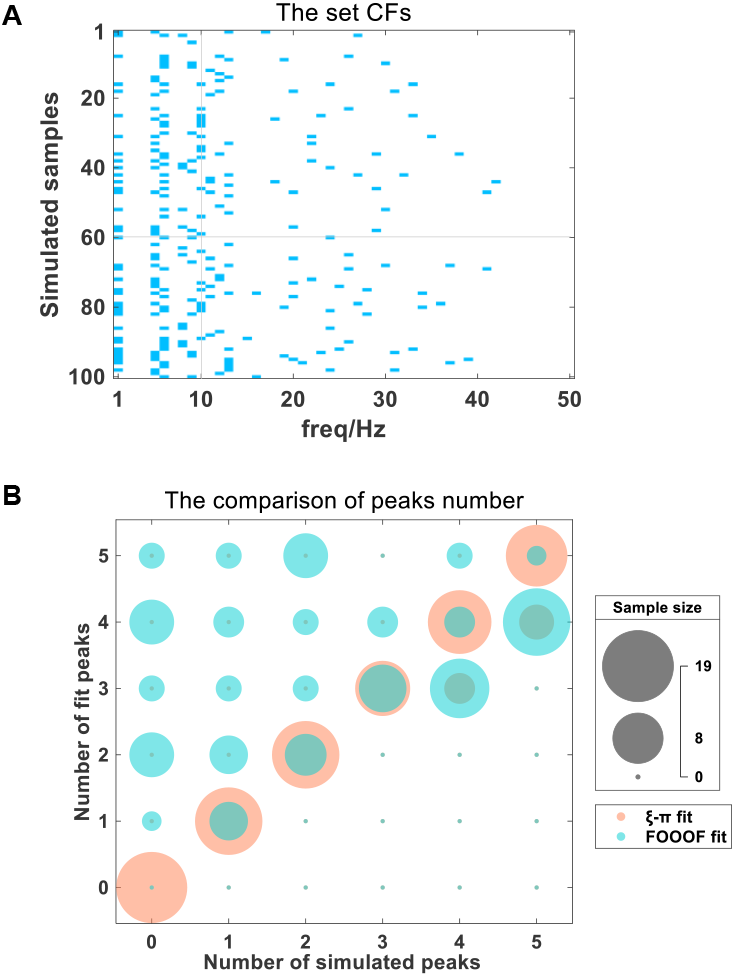
The set CFs and the comparison of peaks number. **(A)**. The set CFs in batch simulation; **(B)**. The number of fit peaks vs. the number of simulated peaks; the bubble radius positively correlates with sample size.

Table. II is the loss statistics of decomposition models for fitting AC/PCs and CF identification. The mean ± deviation of LogMSE in AC (PC) fit was 0.78±1.46 (0.89±1.41) for *ξ*-*π* and 2.98±1.25 (1.49±1.26) for FOOOF. For both AC and PC, *ξ*-*π* generally had much less MSE than FOOOF; especially, the LogMSE increment from *ξ*-*π* to FOOOF in fitting AC was more obvious than that in fitting PC; the LogMSE of *ξ*-*π* did not show much difference between AC fit and PC fit, whereas the LogMSE of FOOOF in AC fit was higher than that of FOOOF in PC fit. This may indicate that *ξ*-*π* is generally better than FOOOF in terms of the fitting error and the stability in AC and PC fit; FOOOF may generally have a larger fitting error than *ξ*-*π* and the AC fit using FOOOF usually produced larger error than the PC fit. Besides, *ξ*-*π* achieved generally correct identification of CFs with accuracy(Acc) 99.86%, precision(Prec) 100%, recall(Rec) 97.15%, and F1 98.56%.

**TABLE II.** Loss statistics of Batch Simulation

| Model | Log(MSE) |  | Identification of CFs |  |  |  |
| --- | --- | --- | --- | --- | --- | --- |
|  | AC | PC | Acc | Prec | Rec | F1 |
| ξ-π | **0.78±1.46** | **0.89±1.41** | **99.86%** | **100%** | **97.15%** | **98.56%** |
| FOOOF | 2.98±1.25 | 1.49±1.26 | 96.68% | 62.05% | 83.74% | 71.28% |

### B. Validation from Real Data

#### 1) Sleep State Classification using CCSHS EEG

Fig. 6A is the decomposed AC and the power law fit to the C4 spectra of subject 1800001 at the wake-N2-REM sleep state. Unlike FOOOF inherently adopted the power law function and obtained the AC exponent simultaneously, *ξ*-*π* only obtained the AC after decomposition. The power law function was applied subsequentially to the *ξ*-*π* decomposed AC so as to obtain the AC exponent. Hence, the left and middle plots displayed the decomposed AC and further power law fit to the *ξ*-*π* decomposed AC correspondingly, whereas the right plot showed the AC directly fit with the power law function by FOOOF. The comparison between the left and the right plots clearly showed that *ξ*-*π* could keep the natural shape characteristic of three sleep states by ignoring the low amplitude loss at high frequencies, while FOOOF only fitted the rough trend with a linear decreasing function in the log-log scale.

**Fig. 6.**
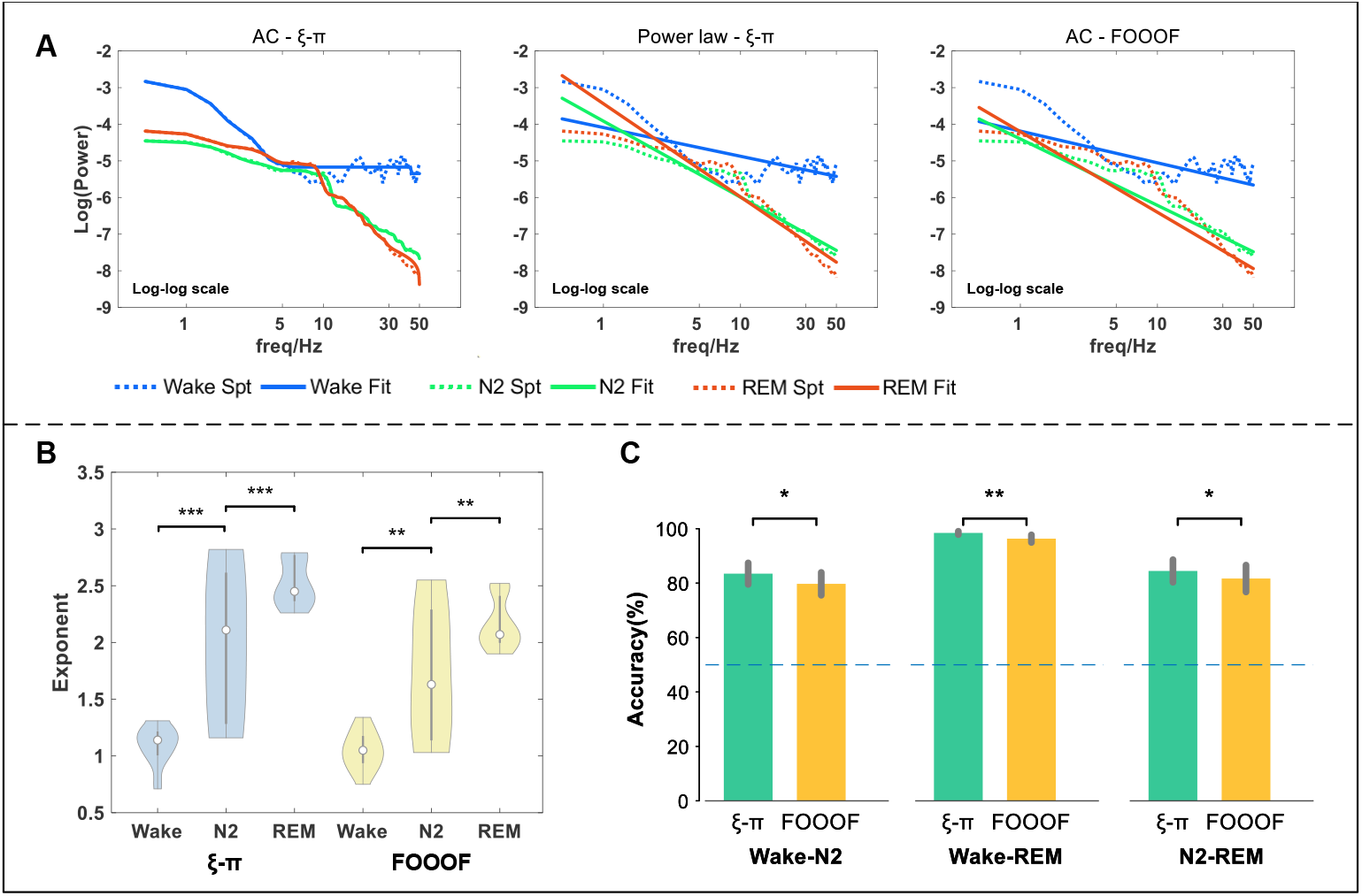
Sleep state classification using AC exponent. **(A)**. An illustration of AC fit to estimate AC exponent in decomposing the Wake-N2-REM spectra. (B). Violin plots of AC exponents under the wake, N2, REM states, and the interstate statistical significances of AC exponents. (C). The prediction accuracies. *: p<0.05, **: p<0.01, ***: p<0.001.

In addition, the comparison between the middle and the right plots indicated that all the included angles of wake-N2, N2-REM, and wake-REM derived from *ξ*-*π* were greater than that from FOOOF, meaning the greater distances of AC exponents within any pair of sleep states. This can be confirmed from that the AC exponents of wake, N2, and REM by *ξ*-*π* and FOOOF were 0.784, 2.077, 2.547, and 0.866, 1.813, 2.199, respectively. Thus, Fig. 6A illustrated that *ξ*-*π* yielded greater interstate differences of AC exponents, resulting from a better fit than FOOOF.

Fig. 6B shows the violin plots of AC exponents of three sleep states after the spectral decomposition was applied. The AC exponents generally showed an increasing trend from wakefulness to N2 and then to REM for both two models. This meant that the AC exponent was positively associated with sleep depth, which was in line with the previous findings [46].

The independent sample t-test was made to measure the interstate difference of the AC exponent, which followed the way of measuring the interclass distance of features in a classification task, reflecting the interclass distinguishability. Based on *ξ*-*π*, the two p values for the pair “wake-N2” and the pair “N2-REM” were *<*0.001 (***), while based on FOOOF, the p values of the pair “wake-N2” and the pair “N2-REM” were *>*0.001 but *<*0.01 (**). This proves that *ξ*-*π* helped improve interstate distinguishability using the AC exponent. *ξ*-*π* may be an effective method in EEG biomarker extraction for sleep staging studies.

Fig. 6C shows the prediction accuracy of the LDA model using AC exponents to discriminate sleep states. In each bar plot, the mean accuracy and the standard deviation were estimated over 19 collections. The mean accuracies of *ξ*-*π* vs. FOOOF were 83.47±17.22% vs. 79.73±18.28% for Wake-N2, 98.43±2.67% vs. 96.33±6.08% for Wake-REM and 84.46±18.11% vs. 81.68±21.48% for N2-REM. The independent sample t-test was performed on the accuracies of 19 collections between *ξ*-*π* and FOOOF. The p values within each pair of “Wake-N2”, “Wake-REM” and “N2-REM” were all *<*0.05. This indicated that the predicting accuracies based on *ξ*-*π* were significantly higher than that based on FOOOF. Besides, the standard deviations of accuracies over 19 collections based on *ξ*-*π* were all less than that based on FOOOF. Hence, Fig. 6C meant that *ξ*-*π* was more stable and significantly more accurate than FOOOF in predicting the sleep states.

#### 2) Peak discovery using MNI iEEG

Fig. 7 illustrates spectral decomposition on an iEEG channel from the middle temporal gyrus randomly picked from the MNI iEEG dataset. Each sub-plot displayed the decomposition with individual components and its “log-log” plot which showed the GF but omitted the PC. The raw spectra mainly contained 1 AC and 1 PC that was slightly skewed and concave upward. *ξ*-*π* decomposed the raw spectra into 1 AC and 1 PC, whereas FOOOF decomposed the raw spectra into 1 AC and 2 PCs.

**Fig. 7.**
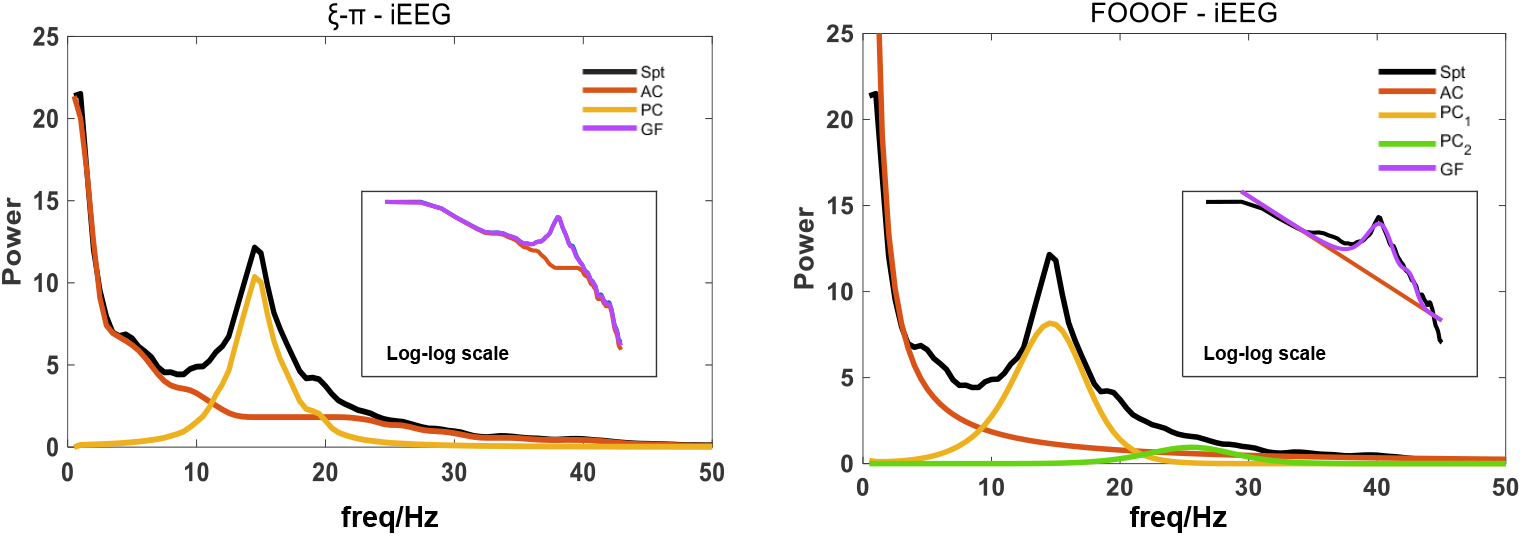
An illustration of spectral decomposition on an iEEG channel from the middle temporal gyrus. Spt: spectra; AC: aperiodic component; PC: periodic component; GF: global fit.

The AC fit by *ξ*-*π* generally followed the monotonically decreasing trend and had well coincidences in the *<*8Hz low frequencies, while the AC fit by FOOOF only roughly captured the real AC trend but missed the fit in low (*<*4Hz) frequencies and the local details at 10-30Hz. The PC fit by *ξ*-*π* was only one component, all under the raw, and nearly kept the peak shape, while the PC fit by FOOOF were two Gaussian curves. The major PC by FOOOF showed an even larger bandwidth and a CF slightly shifted toward high frequency, compared to the raw peak, and the minor PC by FOOOF seemed to overfit in the frequency range where no peak existed most probably. The GF difference between *ξ*-*π* and FOOOF can be inspected from the log-log plot. The GF by *ξ*-*π* generally coincided with the Spt and the tiny fit error in the high frequency *>*20Hz can be neglected. Nevertheless, the GF by FOOOF hardly fitted the Spt. The fitting error of GF by FOOOF would be more apparent after the log scale was back-transformed to the natural scale.

Besides, Table. III shows the comparison quantified by the evaluation metrics with the mean (± standard deviation) over 1772 channels of iEEG. The LogWLS of *ξ*-*π* with -0.8±1.5 were generally much lower than that of FOOOF with 1.9±1.4. The *R*^2^ of *ξ*-*π* with 0.9976±0.0079 was generally higher than that of FOOOF with 0.9867±0.0109. And the Whittle likelihood of *ξ*-*π* with -212.88±182.13 were generally smaller than that of FOOOF with -201.01±117.28. It was found that *ξ*-*π* had a lower Whittle likelihood than FOOOF for 97.2% channels, meaning the superiority of *ξ*-*π*.

**TABLE III.**
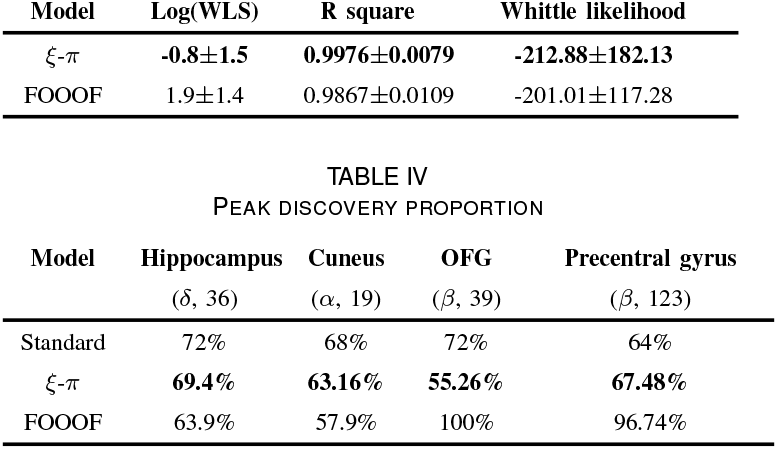
Loss statistics on MNI iEEG

The empirical standard from the resting MNI spectra atlas summarized in the section II-E.2 were filled in the Table.IV together with the reported proportion by decomposition models. For hippocampus (*δ* peak) and cuneus (*α* peak), although both models derived lower proportions than the empirical standard, *ξ*-*π* was closer to the empirical standard with 5.34% increment on average than FOOOF. For the OFG and the precentral gyrus, *ξ*-*π* detected 16.74% less and 3.48% more peaks than the empirical standard, respectively; whereas FOOOF reported the existence of *β* peak in almost all channels, which was far from the standard. Except that *ξ*-*π* found 6 more channels in the OFG (36 channels) presenting *δ* peak than the empirical standard, the peak discovery by *ξ*-*π* nearly approached the clinical finding with a 3.67% deviation on average for the other three brain regions.

Hence, *ξ*-*π* was generally much closer than FOOOF to the empirical standard in peak discovery. FOOOF was not as good as *ξ*-*π* at detecting peaks in the low-frequency range, such as delta and alpha peaks, and much more sensitive than *ξ*-*π* to detect the beta peaks, even if no beta peak existed. The high sensitivity of FOOOF in detecting beta peaks may validate the high OER found in the section III-A.2. Both loss statistics and the validation against the clinical evidence suggest that *ξ*-*π* was effective.

## Iv. Discussion

The advantages of *ξ*-*π* were summarized as follows: **(1) Great robustness to irregular shape**. *ξ*-*π* would perform well by following the natural shape characteristics without the restriction of kernel functions When the spectra have multiscale AC, the AC exponent heavily depends on the frequencies [19], [48], [49], two CFs are close as shown in Fig. 4A [50], or the spectra present other shape irregularities. **(2) Flexible parameterization**. *ξ*-*π* precisely decomposes the spectra as the first step and then offers the chance of flexibly using the descriptive function to users for parameterization [51]. Based on the individual PCs, the amplitude, bandwidth, and CF are easily measurable with descriptive statistics where kernel functions may be unnecessary. **(3) Natural scale**. The natural scale helps display the prominence and predominance of PCs fairly for both low-frequency and high-frequency PCs, refraining the model from being more sensitive to the high-frequency low-amplitude peaks that will be magnified in the log scale [52]. As a counterexample, FOOOF detected peaks that may not exist in the *β* band, as shown in Table. IV. The scale transformation not only matters the visual prominence of PCs but also determines the mixing mechanism. The additive mechanism in the log scale equals to the multiplicative mechanism in the natural scale. **(4) Additive mechanism**. *ξ*-*π* fits in the natural scale using the additive mechanism, helping for interpretation. The additive mechanism is the most parsimonious viable for decoding resting state activity and most event-related activities. The level of 1/f activity and alpha power are not positively correlated within participants, in line with the additive but not the multiplicative mechanism [27]. The multiplicative mechanism should not be used because it involves nonlinear operations and problematic assumptions [27], [53]. The multiplicative mechanism may be only reasonable when the same underlying source in processing event-related brain activities perfectly couples the baseline AC and the narrowband PC caused by experimental stimuli. It is the multiplicative mechanism used in FOOOF that will greatly enlarge the fitting error and interfere with the individual fit.

The utilization of *ξ*-*π* should take care of the following points. **(1) Uncertainty in the spectral estimate**. Depending on the method used to estimate the power spectrum and the amount of data used, there may be spurious peaks resulting from noise, or obscured peaks due to spectral bias. Before the spectral estimation, we recommend the users clean the EEG recordings by manually selecting the quasi-stationary noise-free epochs or adopting the advanced preprocessing tools [54], [55] and quality control measures as we proposed in [56], [57]. The Welch’s window smoothing method, the mutitapers approach, and the autoregressive model can make the spectrum smooth [58]–[60]. Since the multitapers approach is a frequency-specific method and not sensitive to weak noise, it can be a good choice to characterize the neural oscillations [61]. **(2) Non-sinusoidal neural oscillation**. The sinusoidal assumption in the Fourier spectra may generate spurious PCs because of the raise-decay asymmetry of non-sinusoidal oscillations [28]. Non-sinusoidal periodic signals may have various waveform shapes that are ununimodal [62]– [64]. Non-sinusoidal waveforms resulted in the harmonics with multiple PCs appearing in the spectral curve. *ξ*-*π* will identify all PCs without bias to any of them purely based on thresholds in the “PC detection” step. After decomposition, the harmonics can be determined by: (a) CFs are equidistant, such as *f*_0_, 2*f*_0_, 3*f*_0_, 4*f*_0_; (b) the maximum amplitudes of PCs decrease with the CFs getting larger. The existence and the operation of harmonics come to the individual PCs. For instance, the interpretation of PCs can be mainly ascribed to the fundamental PC at *f*_0_. Since the nonsinusoidal waveforms are stereotyped, *ξ*-*π* can be extended with a ‘directory’ of non-sinusoidal basis functions, the matching pursuit algorithm, and the empirical mode decomposition and applied together with the evaluation and measurement of waveform and cycle properties [63]. The cross-frequency coupling analysis may help disassociate the harmonics and the non-harmonics [65]. **(3) “AC knee vs weak PC”** The following two situations may be obscured to lower the identifiability of AC and PCs. Weak PC: the AC slope obscures the left side of a weak PC that doesn’t clearly show a local maximum, making the left side of PC appear as a flattening of the AC slope; AC knee: the AC may have a lower slope to the left of a “spectral knee” and a higher slope to the right [23], [48], [66]. We have made detailed simulations of these two situations and summarized the following findings. It is indeed possible that the weak PC and AC knee compromise each other. However, the discussion about the AC knee and weak PC that interferes with the AC slope only makes sense in the log-log scale. *ξ*-*π* doesn’t work with the log-log scale but only performs the fitting on the natural scale. As the first step of *ξ*-*π*, the “PC detection” is the key to its identifiability of AC and PCs. The “PC detection” mainly adopts the MATLAB “findpeaks” routine, which identifies peaks by a few preset thresholding parameters. In the case of weak PC, the weak PC becomes more prominent in the natural linear-linear scale than that in the log-log scale. If the local maxima of a weak PC can be detected by “findpeaks”, then *ξ*-*π* can identify the PC and finally output the PC fit. Otherwise, if the maxima of weak PC don’t exist in the natural scale after being mixed with AC, it becomes impossible for “findpeaks” to identify a local maximum even though the thresholding parameters can be finely tuned. In the case of AC knee, the spectral curve is only AC with varying slopes and knee. No matter in the log-log scale or the natural scale, it is a monotonically decreasing function. The *ξ*-*π* model will fit the AC knee with a monotonically decreasing function based on the shape language modeling (SLM). In the case of “AC knee vs. weak PC”, the identifiability of *ξ*-*π* model relies on (a) the existence of a local PC maximum; (b) whether the local PC maximum can be found by the “findpeaks” function in the “PC detection” step. *ξ*-*π* is currently a data-driven dominant approach based on the shape characteristics. In the future, the determination of AC knee without a local maximum can be taken as a prior that *ξ*-*π* should fit a minor PC around the knee location in the natural scale. More importantly, *ξ*-*π* can be updated to a biophysically model-driven approach by incorporating more neurophysiological priors relating to neural oscillation, such as the synaptic firing mechanism, the neural field theory, and the volume conduction model.

The limitations and outlook are: (1) *ξ*-*π* can be extended with the brain connectivity, the electrophysiology source imaging [67], and the dynamic spectra decomposition [21]. (2) As FOOOF is the most recent, popular, and typical parametric model, it was taken as the main competitor to compare with *ξ*-*π*. (3) Note that the main goal of this paper was to propose the nonparametric model *ξ*-*π* and demonstrate its superiority in decomposition and fitting. The parameterization on the AC exponent and the CFs of PC provided additional supportive evidence. Complete investigation on parameterization is beyond the current scope, whose worth will be excavated through multiple quantitative neurophysiology studies in the future.

## V. Conclusion

This study presented *ξ*-*π* as a nonparametric model to address the issues in dealing with shape irregularities, statistical distribution, and scale transformation and to resolve the spectral mixing problem of multiple neural oscillatory processes. Its advantages were demonstrated with elaborated simulation, together with the sleep state classification and the peak discovery studies through neurophysiological evidence. Its applications may be extensively explored in cognitive neuroscience, mental disorders, brain-machine interface, etc. The MATLAB packages and tutorials are freely available at https://github.com/ShiangHu/Xi-Pi.

